# Guanylate cyclase activity of TIR1/AFB auxin receptors mediates rapid auxin responses

**DOI:** 10.1101/2023.11.18.567481

**Authors:** Linlin Qi, Huawei Zhai, Huihuang Chen, Mateusz Kwiatkowski, Jieren Liao, Mengting Lu, Yuanrong Pei, Yongqiang Gao, Ping Yun, Lanxin Li, Sergey Shabala, Edward E. Farmer, Krzysztof Jaworski, Jiří Friml

## Abstract

The plant hormone auxin is perceived by TIR1/AFB receptors, which mediate transcriptional reprogramming via a ubiquitination- and cAMP-dependent mechanism. Same receptors mediate also rapid, non-transcriptional growth responses that are mechanistically less well understood. Here, we demonstrate that TIR1/AFB receptors possess not only adenylate cyclase but also guanylate cyclase activity. Auxin perception independently activates cAMP and cGMP production by TIR1/AFBs *in vitro* and elevates both cAMP and cGMP levels *in planta*. Genetic and synthetic biology approaches revealed that AFB1 guanylate cyclase activity is required for the rapid cytosolic Ca^2+^ transients and root growth inhibition. This unprecedented combination of adenylate and guanylate cyclase activities within a single hormone receptor establishes a new paradigm for how one perception mechanism can mediate distinct downstream responses.

**One-Sentence Summary:** TIR1/AFB auxin receptors possess both, adenylate and guanylate cyclase activities, mediating transcriptional reprogramming and rapid Ca^2+^-dependent growth responses, respectively.

## Main Text

Auxin is one of the most prominent plant hormones and much of its action on growth and development has been attributed to its regulation of transcription (*1, 2*). The TIR1/AFB auxin receptors are F-box subunits of the E3 ubiquitin ligase complex, which after auxin perception associates with the Aux/IAAs co-receptors leading to the ubiquitination and degradation of the latter. Since Aux/IAAs act also as transcriptional repressors, by interacting with and inhibiting ARF transcription factors, their auxin-triggered degradation contributes to ARFs activation to promote transcription of auxin-regulated genes (*3-6*). This established canonical auxin signaling pathway explains most of the auxin developmental roles. Recent accumulating evidence suggests that TIR1/AFBs mediate also rapid, non-transcriptional cellular responses in root elongation regions, primarily cytosolic Ca^2+^ transients acting upstream of apoplastic alkalinization and membrane depolarization; all crucial for rapid, auxin-induced root growth inhibition and root gravitropism(*7-14*).

Recently, adenylate cyclase activity (AC) has been identified in TIR1/AFB auxin receptors, which generates cAMP after auxin perception, however, this activity is not required for rapid responses but for transcriptional regulations (*15, 16*). Thus, mechanism underlying rapid auxin responses remains elusive.

Here we show that TIR1/AFB auxin receptors, next to AC, also have guanylate cyclase (GC) activity, which is stimulated by auxin perception and produces another important second messenger, cGMP. In contrast to AC, the GC activity is required for the rapid non-transcriptional responses including cytosolic Ca^2+^ transients, suggesting cGMP mediates rapid cellular responses converging on root growth regulation.

## Results

### TIR1/AFBs from angiosperms have guanylate cyclase activity

The AC activity of TIR1/AFBs is not essential for the rapid non-transcriptional auxin responses (*15*), thus prompting us to search for additional TIR1/AFB functionalities. Besides cAMP, a similar small molecule cGMP, the product of GC activity, is another key second messenger in animals (*17*). By a scrutinized sequence analysis, we found a relatively conserved GC motif (*18*) adjacent to the previously characterized AC motif (*15*) in the C-terminal region of TIR1/AFBs (Fig. 1A). To test the potential GC activity of TIR1/AFB auxin receptors, we used the His-GFP-Flag-TIR1 purified from Sf9 insect cells, and GST-AFB1 as well as GST-AFB5 proteins purified from *E. coli* (*15*). We performed the *in vitro* GC activity assay, followed by cGMP detection using liquid chromatography–mass spectrometry (LC–MS/MS) method. cGMP was reliably detected in all reactions with different TIR1/AFB proteins tested (Fig. 1B and fig. S1) and typical Michaelis-Menten kinetics were obtained (Fig. 1, C to E) with parameters of the maximum velocity (V_max_) and the Michaelis constant (K_M_) similar to other reported plant GCs (*19-23*). The GC motif is less conserved compared to the AC motif and it was originally identified only in TIR1 and AFB1sequences. The confirmation of the GC activity also for AFB5 indicated that the originally reported GC motif could be more relaxed.

**Fig. 1.**
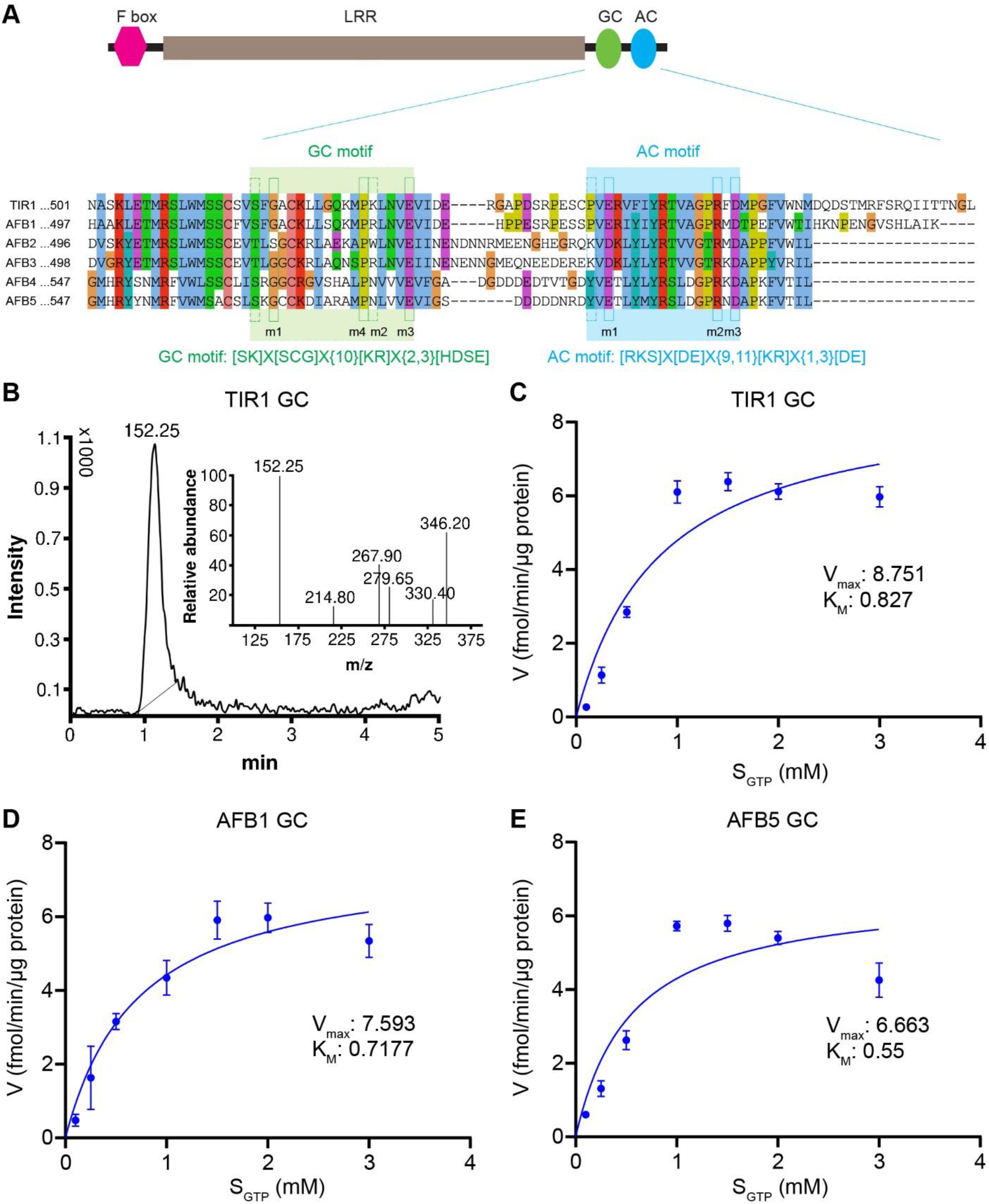
TIR1/AFB auxin receptors have GC activity. (**A**) Simplified scheme showing the domain structure of TIR1/AFB auxin receptors and a multiple sequence alignment showing the conserved GC and AC motif in the C-terminal region. LRR, leucine-rich repeat. m1, m2, m3 and m4 are conserved key amino acids that were mutated to alanine to disrupt the GC or AC activity in subsequent experiments. (**B**) A representative LC-MS/MS spectrum showing cGMP detection in the GC reaction of His–GFP–Flag–TIR1, with the characteristic peak used for quantification. (**C**) GC activity of His–GFP–Flag–TIR1. (**D**) GC activity of GST-AFB1. (**E**) GC activity of GST-AFB5. His–GFP–Flag–TIR1 was purified from Sf9 insect cells. GST-AFB1 and GST-AFB5 were purified from *E. coli. In vitro* GC activity assays were performed with the purified proteins followed by cGMP quantification using LC-MS/MS. Michaelis–Menten kinetics were generated using non-linear regression in GraphPad Prism 8. S_ATP_, substrate (ATP) concentration; V, velocity. Data are mean ± s.d. of three biological replicates.

To evaluate the evolutionary conservation of the TIR1/AFBs GC activity, we compared the TIR1/AFB orthologous sequences from different representative species during the evolution of land plants. There are four key amino acids (a1-a4) in the GC motif (fig. S2A). Whereas a1 and a4 are highly conserved across all the orthologous sequences analyzed, a3 is divergent everywhere. The a2 residue diverged before angiosperms, and became fixed in the angiosperm GC motif, suggesting that GC activity may have evolved only in this lineage. To test this hypothesis, we purified the GST-PaAFB1 and GST-PaAFB2, which are the only two closest TIR1/AFB orthologues from the gymnosperm *Picea abies*, and GST-AmTrAFB2 and GST-AmTrAFB3 which are two out of the three TIR1/AFB orthologues from the ancestral angiosperm *Amborella trichopoda* (fig. S3A), and tested their potential AC and GC activities. As expected, all of them had AC activity (fig. S2B), indicating the proteins purified are biochemically functional. In contrast, only AmTrAFB2 showed clear GC activity (fig. S2C), suggesting that GC activity of TIR1/AFBs indeed evolved in the earliest angiosperm.

Hence, the TIR1/AFB orthologues presumably in most, if not all flowering plant species, may have GC activity.

### GC and AC activities of TIR1/AFBs are independent

The GC and AC motifs are adjacent in sequence and also spatially close to each other in the protein structure (fig. S4). To validate the importance of the C-terminal GC motif for the GC activity of TIR1/AFBs and further test the specificity of the GC and AC motifs for their corresponding activities, we mutated separately three key residues in the C-terminal GC motif of AFB5 (GCm1, GCm2, GCm3 in Fig. 1A) to alanine. Similar mutations had already been generated for the AC motif of AFB5 (ACm1, ACm2, ACm3) (*15*). We purified all the mutated variants of GST-AFB5 (fig. S3B), and performed the *in vitro* GC and AC activity assays.

Clearly, GCms but not ACms decreased the GC activity of AFB5 (fig. S5A). Similarly, ACms abolished the AC activity of AFB5 as shown before (*15*), whereas GCms did not show any significant effects on it (fig. S5B). In other words, GCms specifically interfered with the GC activity but did not affect the AC activity, and vice versa for ACms. To further confirm this, we utilized a previously characterized N-terminal deleted TIR1 version - TIR1ΔNT (*15*), and purified the similarly mutated proteins (fig. S3C). Again, all three GCms specifically interfered with the GC activity of TIR1ΔNT, while all three ACms specifically abolished the AC activity (Fig. 2, A and B). Similar results were obtained for the AFB1^GCm1^ and AFB1^ACm1^ (Fig. 2, C and D, and fig. S3D). Taken together, these results clearly demonstrate that the C-terminal GC motif within TIR1/AFBs is responsible for the GC activity and that GC and AC activities are independent.

**Fig. 2.**
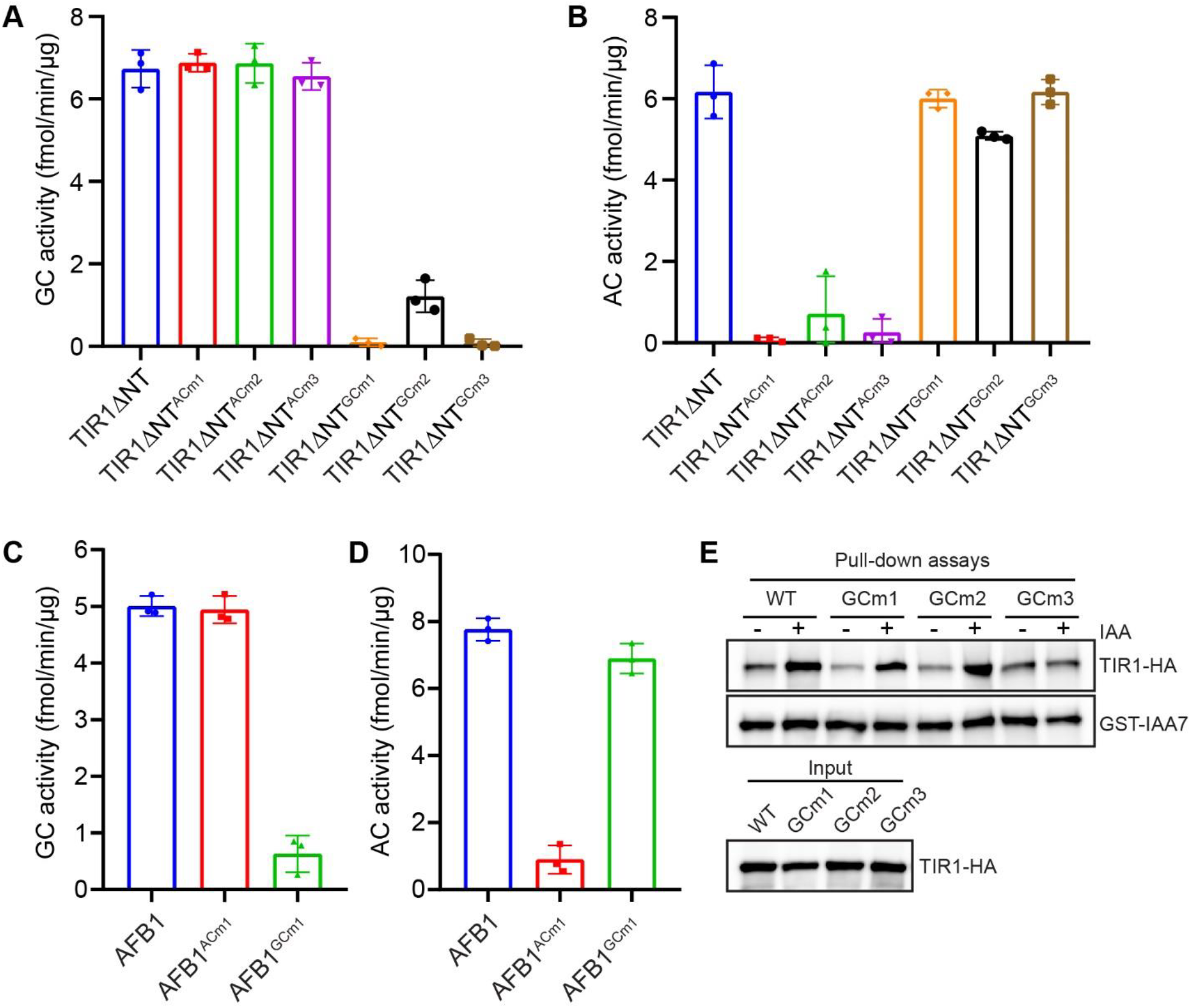
The C-terminal GC and AC motifs are specifically responsible for the GC or AC activity of TIR1/AFBs respectively. (**A**) The C-terminal GC motif but not AC motif is essential and specific for the GC activity of TIR1ΔNT. (**B**) The C-terminal AC motif but not GC motif is essential and specific for the AC activity of TIR1-ΔNT. (**C**) The GC motif is specifically essential for the GC activity of AFB1. (**D**) The AC motif is specifically essential for the AC activity of AFB1. The indicated GST-tagged proteins were purified from *E. coli. In vitro* GC/AC activity assay was performed and cGMP/cAMP was quantified by LC-MS/MS. Data are mean ± s.d. of three biological replicates. Data for the AC activity of TIR1ΔNT, TIR1ΔNT^ACms^, AFB1 and AFB1^ACm1^ have been shown previously (*15*). (**E**) Pull-down assay showing differential effects of TIR1^GCms^ mutations on the IAA-induced TIR1-Aux/IAA interaction. The TIR1-HA and its GCms variants were translated *in vitro* using wheat germ extracts. GST-IAA7 was purified from *E. coli*. Pull-down assays were performed in the presence or absence of 10 μM IAA as indicated. The experiment was repeated three times with similar results.

It had been shown before that one of the ACms — ACm2 also interferes with auxin perception — the auxin-induced TIR1/AFB interaction with Aux/IAAs (*15*). To clarify the similar problem for GCms, we introduced the three GCms into full length TIR1, and performed the pull-down experiments with *in vitro*-translated TIR1-HA variants and purified GST-IAA7 in the presence of IAA. The results showed that whereas GCm3 abolished the auxin-induced interaction between TIR1 and IAA7, GCm1 and GCm2 did not have any visible effects (Fig. 2E, and fig. S6), indicating these two mutations can be used to decrease specifically the GC activity without interfering with AC activity or Aux/IAA interaction.

Overall, this shows that TIR1/AFB auxin receptors have independent AC and GC activities. Thus, TIR1/AFBs represent, to our knowledge, the first identified example of protein combining these two enzymatic activities.

### Auxin perception rapidly stimulates the GC activity

Next, we tested an impact of auxin perception on TIR1/AFB GC activity. We used the His-GFP-Flag-TIR1 protein purified from Sf9 insect cells, and performed the *in vitro* GC activity assay in the presence of auxin (IAA) with or without IAA7 or IAA17 co-receptors. IAA or IAA7/17 alone did not have any significant effect on the TIR1 GC activity, but auxin together with IAA7 or IAA17 clearly enhanced the GC activity (Fig. 3A). To confirm this observation, we performed similar experiments for AFB1 purified from *E. coli* with a similar result (Fig. 3B). Together, this shows that auxin perception enhances the GC activity of TIR1/AFBs *in vitro*.

**Fig. 3.**
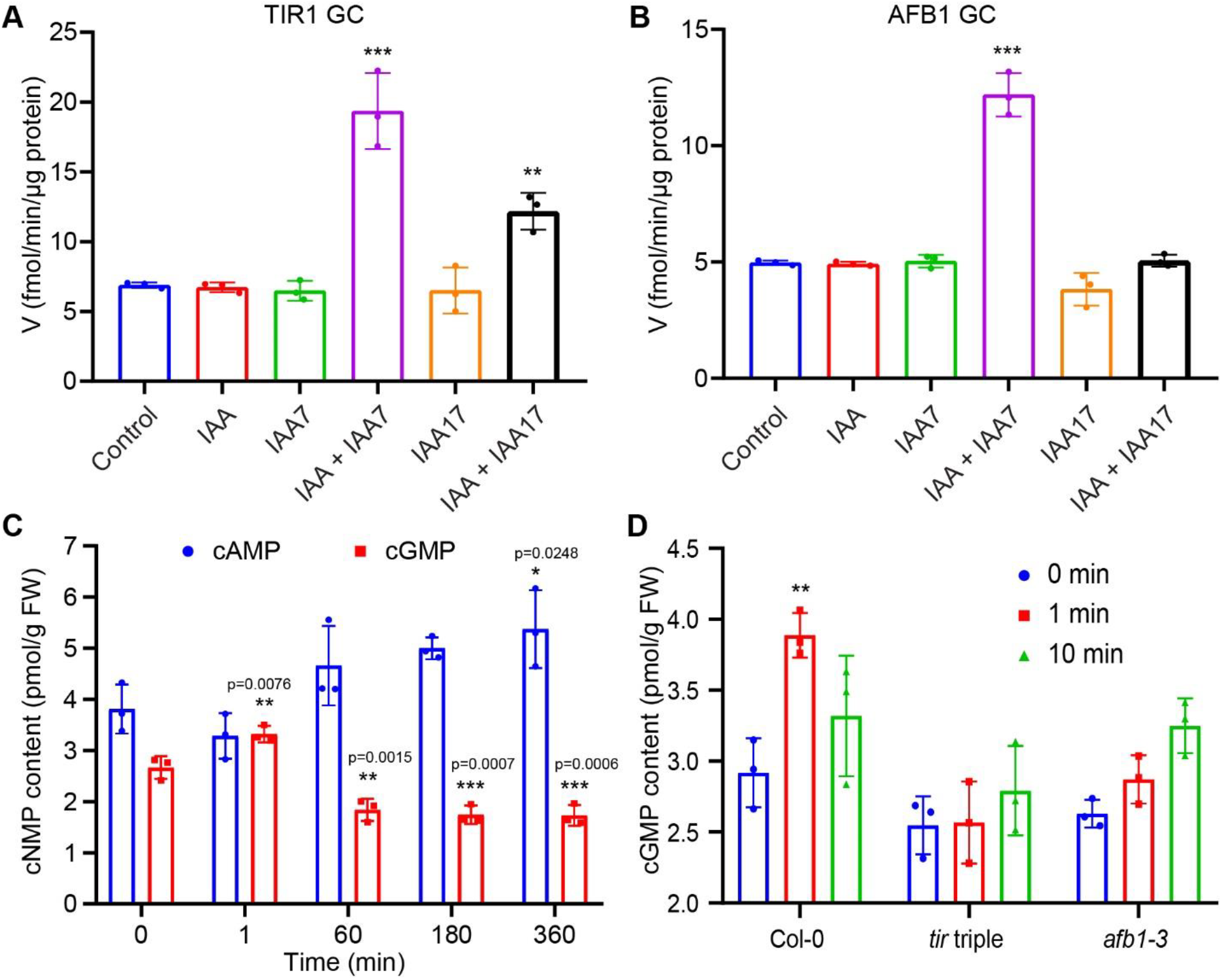
Auxin perception rapidly stimulates the GC activity of TIR1/AFB receptors. (**A**) Auxin together with Aux/IAA stimulates the GC activity of TIR1. (**B**) Auxin together with Aux/IAA stimulates the GC activity of AFB1. *In vitro* GC activity assay was performed for His– GFP–Flag–TIR1 and GST tag-cleaved AFB1 respectively, in the presence of 10 μM IAA, 5 μg IAA7, 5 μg IAA17, and the indicated combinations. cAMP was quantified by LC–MS/MS after the reactions. Data are mean ± s.d. of three biological replicates. One-way ANOVA and Dunnett’s multiple comparisons test. ** *p* ≤ 0.01 (*p* = 0.0032); *** *p* ≤ 0.001. (**C**) Differential dynamics of cAMP and cGMP content in root tissues after auxin treatment. Data for cAMP content have been published previously (*15*), and is shown here for comparison. One-way ANOVA and Dunnett’s multiple comparisons test. Asterisks indicate significant difference between the corresponding group and the control (0 min). (**D**) The auxin-induced rapid increase of cGMP levels in roots is dependent on TIR1/AFB receptors. Two-way ANOVA and Tukey’s multiple comparisons test. ** *p* ≤ 0.01 (p = 0.004). Data are mean ± s.d. of three biological replicates.

Next, we tested how auxin regulates the GC activity *in planta*. We treated Arabidopsis seedlings with 100 nM IAA and harvested root tissues at different time points to measure cAMP and cGMP in the same samples using LC-MS/MS. In contrast to the cAMP dynamics, which showed a slight decrease at 1 min and then steady increase after 1 h (*15*), cGMP level rapidly increased at 1 min after auxin treatment and then decreased after 1 h (Fig. 3C). The differential dynamics of cAMP and cGMP after auxin treatment implies that they may mediate different downstream processes. Similar experiments in the *tir1-1 afb2-1 afb3-1* (*tir* triple) and *afb1-3* mutants showed that the auxin-induced rapid increase of cGMP was almost completely abolished in *tir* triple and largely reduced in *afb1-3* mutants (Fig. 3D). This is consistent with both, *tir* triple and *afb1-3* being defective in rapid auxin growth response (*10, 13, 24*).

These data show that auxin perception stimulates cGMP production by TIR1/AFB receptors *in vitro* and auxin treatment rapidly increases the cGMP levels in roots via the TIR1/AFB receptors.

### AFB1 GC motif is crucial for rapid auxin responses in roots

As the AC activity of TIR1/AFBs is required for transcriptional but not for the rapid, non-transcriptional responses (*15*), we tested whether it may be the GC activity, which is important. It has been established that among the 6 TIR1/AFBs auxin receptors in *Arabidopsis*, the predominantly cytosol-localized AFB1 plays major roles in rapid auxin responses as *afb1* mutant is insensitive to auxin in terms of cellular effects including cytosolic Ca^2+^ transient, apoplast alkalinization and membrane depolarization; processes converging on rapid root growth inhibition (*13, 14, 24-26*). We confirmed these observations in *afb1-3* roots by both, microfluidics and electrophysiology (fig. S7).

To test the importance of the GC motif in AFB1 for its ability to mediate rapid auxin responses, we introduced the GCm1 mutation into AFB1 coding sequence and transformed the construct into *afb1-3* mutant generating *AFB1::AFB1*^*GCm1*^ lines. Nonetheless, AFB1^GCm1^ complemented rapid auxin-induced root growth inhibition of the *afb1-3* mutant to the same extent as wild type (WT) version of AFB1 (AFB1^WT^) (fig. S8A). Measurements of cGMP levels in the root tips revealed that the AFB1^GCm1^-complemented lines were not strongly compromised in the auxin-induced rapid increase of cGMP (fig. S8B). Hence, we designed a GCm1,2,4 higher-order mutation, in which three amino acids in the GC motif (shown in Fig. 1A) were all mutated to alanine and generated the *AFB1::AFB1*^*GCm1,2,4*^; *afb1-3* lines. This mutation fully abolished rapid cGMP increase after auxin treatment (fig. S8B). Accordingly, the auxin insensitivity of root growth in *afb1-3* was not complemented (Fig. 4A), as reproducibly observed in 6 independent *AFB1::AFB1*^*GCm1,2,4*^; *afb1-3* lines (fig. S8C). This suggests that AFB1 GC activity is required for rapid auxin-triggered root growth inhibition.

**Fig. 4.**
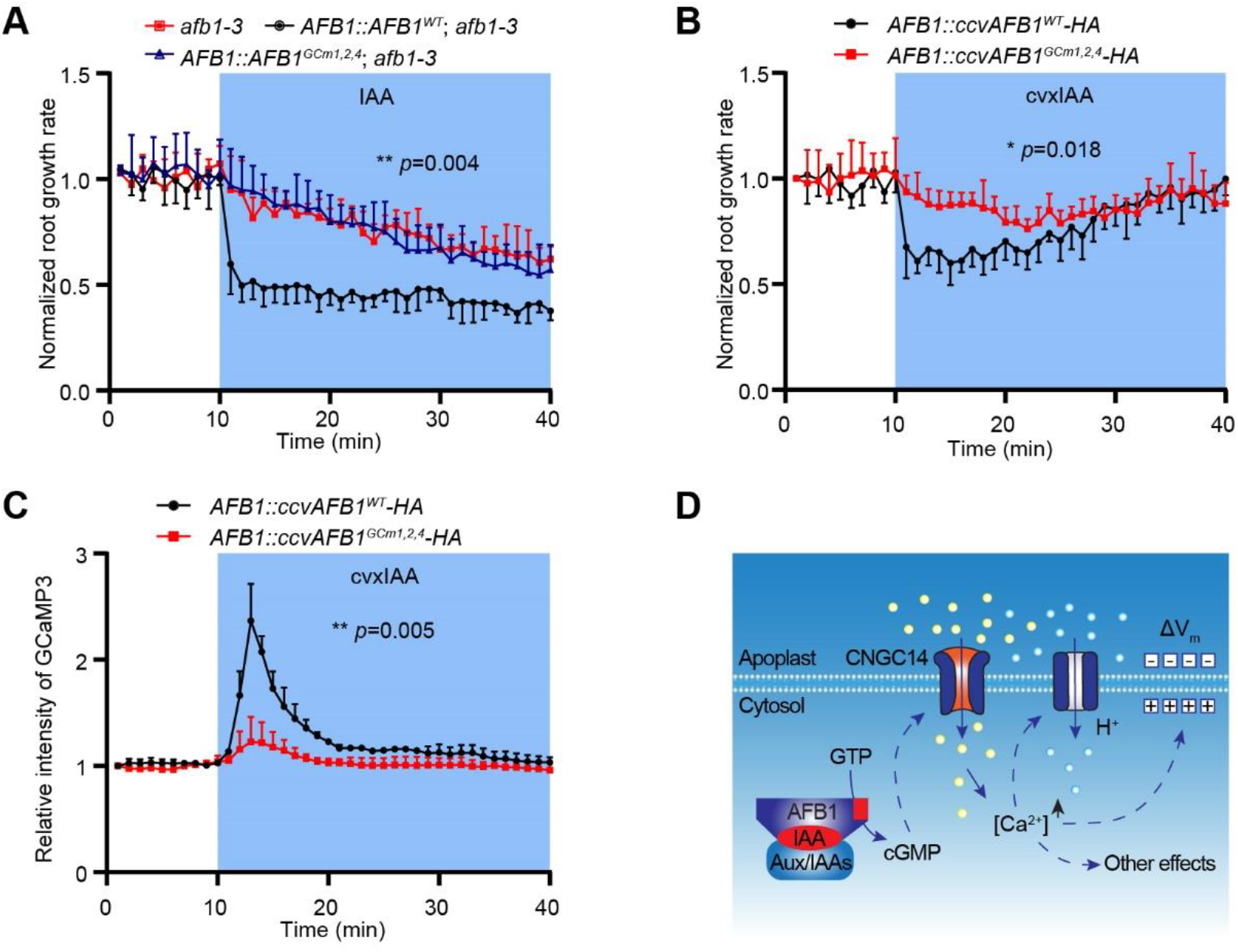
AFB1 GC activity is crucial for rapid auxin responses. (**A**) AFB1^WT^ complements, whereas AFB1^GCm1,2,4^ fails to complement the *afb1-3* mutant in terms of IAA-induced rapid root growth inhibition. (**B**) cvxIAA-induced rapid root growth inhibition is largely reduced in ccvAFB1^GCm1,2,4^ comparing to ccvAFB1^WT^. (**C**) cvxIAA-induced cytosolic Ca^2+^ transients are largely abolished in ccvAFB1^GCm1,2,4^ comparing to ccvAFB1^WT^. The vRootchip experiments were performed with the indicated genotypes. Mock medium was changed to medium containing 10 nM IAA or 50 nM cvxIAA at 10 min time point. Normalized root growth rate and relative fluorescence intensity of GCaMP3 in elongation zone were quantified. Data shown are means +/- s.d. n = 3 or 4 seedlings. Two-way ANOVA and Sidak’s multiple comparisons test. The *p*-values for genotypic variance are shown. (**D**) Schematic diagram showing the rapid non-transcriptional auxin signaling axis: AFB1 – cGMP – CNGC14 – Ca^2+^ acting upstream of apoplast alkalinization, membrane depolarization converging on root growth inhibition.

To further test this, we adapted the cvx/ccv orthogonal synthetic system (*27*) enabling specific activation of AFB1 with a minimal interference from the other TIR1/AFBs (*26*). To this end, we generated ccvAFB1, which does not bind IAA but its synthetic analogue cvxIAA. We prepared the *AFB1::ccvAFB1-HA* and introduced the similar GC mutations into this construct. The obtained constructs were transformed into the background of GCaMP sensor (*28*) for the cytosolic calcium monitoring. As expected, in the *ccvAFB1*^*WT*^ transgenic lines, cvxIAA consistently triggered rapid root growth inhibition and calcium transients (fig. S9, A and B) and similar outcome was observed in *ccvAFB1*^*GCm1*^ line (fig. S9, C and D). In contrast, in the *ccvAFB1*^*GCm1,2,4*^ line, the cvxIAA-induced rapid root growth inhibition and calcium transients were severely compromised (Fig. 4, B and C). Importantly, the defect of ccvAFB1^GCm1,2,4^ in mediating rapid Ca^2+^ transients was observed across 8 independent transgenic lines (fig. S8D), with comparable protein levels as detected in the *ccvAFB1*^*WT*^ and *ccvAFB1*^*GCm1,2,4*^ lines (fig. S8E) and the GCm1,2,4 mutation did not affect the cvxIAA-induced interaction of ccvAFB1 with IAA7 (fig. S8F).

Taken together, these data demonstrate that the GCm1,2,4 mutation in AFB1 abolishes the auxin-induced, AFB1-mediated cGMP elevation *in planta* without affecting its auxin perception function, and renders AFB1 unable to mediate the rapid auxin responses. This strongly suggests that cGMP production by AFB1 is required for rapid auxin-triggered cytosolic Ca^2+^ transients and root growth inhibition.

## Discussion

### AFB1-cGMP-CNGC14 signaling mechanism for rapid responses

TIR1/AFB auxin receptors have been typically associated with transcriptional reprogramming (*6*). However, lately it became clear that they can mediate also rapid cellular responses, including cytosolic Ca^2+^ spikes (*9*), apoplastic alkalinization (*12*) and plasma membrane depolarization (*13*); all linked to rapid, auxin-induced root growth inhibition and root gravitropism (*10*).

Downstream of the receptors, the CNGC14 channel had been demonstrated to be the key component of this regulation (*8*). However, how TIR/AFBs as F-box proteins mediate those cellular responses has been a mystery and remained so also after the identification of the TIR1/AFBs AC activity since the cAMP generated downstream of auxin perception is critical for the transcriptional responses (*16*) but not for the rapid root responses (*15*).

Here we show that TIR1/AFBs have also GC activity and that this feature of TIR1/AFBs is required for the rapid auxin responses including cytosolic Ca^2+^ increase and root growth inhibition. First, auxin treatment rapidly and transiently increases cGMP accumulation in roots (Fig. 3, C and D, and fig.S8B), in a similar temporal pattern as those rapid auxin responses; Second, auxin-induced cytosolic Ca^2+^ spike and rapid root growth inhibition, mediated by the predominantly cytosolic localized AFB1, requires specifically the GC activity of AFB1 (Fig. 4, fig. S7 and fig. S8). These observations imply a linear AFB1 – cGMP - CNGC14 - Ca^2+^ non-transcriptional signaling pathway (Fig. 4D), providing a plausible mechanism for the rapid auxin responses.

Since animal homologues of plant CNGC channels are well-known to be regulated by cyclic nucleotides, there has been an ongoing debate whether this applies also to plant CNGCs (*29*). After recent, structural insights, it seems that plant CNGCs are unable to bind directly cyclic nucleotides (*30*). Nonetheless, our results clearly show that AFB1 can trigger CNGC14-mediated Ca^2+^ transients only if it is able to generate cGMP following auxin perception. This implies that at least CNGC14 can be activated by cGMP, whether this happens directly or indirectly or by which mechanism, remains a topic for future investigations.

### Tri-functional TIR1/AFB auxin receptors

The classical models of the TIR1/AFBs-mediated signaling mechanism has assumed the degradation of Aux/IAA repressors following TIR1/AFB-mediated ubiquitination as the only signaling output from the TIR1/AFB receptors. Now with a recent discovery of their AC (*15*) and GC activities, we show that TIR1/AFBs have three separate functionalities: E3 ubiquitin ligase as well as AC and GC. Notably, those activities can be mapped to distinct domains within TIR1/AFB proteins and can be abolished by specific mutations, showing that they are independent (Fig. 2 and fig. S5). *In vitro*, auxin perception enhances the AC and GC activities and in planta, auxin also stimulates cAMP and cGMP accumulation (*15*) (Fig. 3 and fig. S8B). Hence, following auxin perception, TIR1/AFBs receptors directly generate three independent signaling outputs: Aux/IAA degradation, cAMP and cGMP production. While Aux/IAA degradation and cAMP contribute to transcriptional regulation (*15, 16*), cGMP is crucial to mediate the rapid responses (Fig. 4 and fig. S8).

Such a multiple functionality at the level of receptors is unique throughout the eukaryotic kingdoms and provides a mechanical explanation how the same signal – auxin – can induce such a variety of diverse rapid and sustained cellular and developmental responses.

### Potential role of cGMP as second messenger in plants

Similar to cAMP, cGMP is also a well-known second messenger in mammalian research, acting in peptide and NO signaling (*17, 31*). However, whether cGMP acts as second messenger in plants has remained controversial, even though the list of plant proteins potentially having GC activity *in vitro* keeps growing in recent years (*18, 32*) and includes prominent candidates such as brassinosteroid and phytosulfokine receptors (*21, 23*). This is mainly due to the lack of convincing genetic evidence and failure in identifying the downstream effectors. The identification of the GC activity in TIR1/AFBs receptors with CNGC14 as a likely downstream cGMP target provides a new avenue into studies of cGMP as a second messenger in plants.

In animal cells, cAMP and cGMP are ubiquitously present second messengers involved in dozens of different signaling pathways. They are produced separately by specialized enzymes downstream of distinct signal perception mechanisms (*17*). In contrast, in plants, TIR1/AFBs receptors directly produce in parallel both cAMP and cGMP by independent enzymatic activities to mediate different subsets of responses, presumably via distinct downstream targets. This makes TIR1/AFBs the only known example in any organism of a protein combining both AC and GC activities allowing this auxin pathway to mediate broad range of responses so typical for auxin. Thus, this work contributes to establishing the roles of cAMP and cGMP as second messengers in plant signaling and highlights the unique, plant-specific aspects of their production and action as compared to other eukaryotes.

## Acknowledgments

This research was supported by the Lab Support Facility (LSF), the Imaging & Optics Facility (IOF) and the Nanofabrication Facility (NFF) of IST Austria (ISTA). We are grateful for the seeds from Mark Estelle.

## Funding

European Research Council Advanced Grant ETAP-742985 (JF)

Austrian Science Fund (FWF) I 6123-B (JF)

National Natural Science Foundation of China (32470327) (LQ)

Yuelushan Laboratory Talent Program (2025RC2005) (HC)

## Author contributions

Conceptualization: LQ, JF

Methodology: LQ, MK, KJ, HZ, IK, CH, LL, JL, YG, EEF, PY, SS

Investigation: LQ, MK, HZ, IK, CH, YG, PY

Visualization: LQ, MK, HZ, IK, CH, YG, PY

Funding acquisition: JF, LQ

Project administration: LQ, JF

Supervision: JF, KJ, EEF, SS

Writing – original draft: LQ, JF

Writing – review & editing: LQ, JF

## Data and materials availability

All data are available in the main text or the supplementary materials.

## Supplementary Materials

Materials and Methods

Figs. S1 to S9

Table S1

