## Supplementary Materials for "Guanylate cyclase activity of TIR1/AFB auxin receptors mediates rapid auxin responses"

**The PDF file includes:**

Materials and Methods
Figs. S1 to S9
Table S1

### Materials and Methods

Plant materials and growth conditions

All of the *Arabidopsis* mutants and transgenic lines used in this study are in the Columbia-0 (Col-0) background. The *tir1-1 afb2-1 afb3-1* (*33*) and *afb1-3* (*24*) mutants have been described previously. The calcium sensor GCaMP3 (*28*) was used. To generate the transgenic lines *AFB1::AFB1, AFB1::AFB1*$^{GCms}$ *in afb1-3,* and *AFB1::ccvAFB1-HA, AFB1::ccvAFB1*$^{GCms}$*-HA* in Col-0 or GCaMP3, the MultiSite Gateway Three-Fragment Vector Construction Kit (Life technologies, 12537-023) was used. AFB1 promoter was cloned into pDONR P4-P1r, AFB1 CDS was cloned into pDONR221, AFB1 CDS without stop codon has already been cloned into pENTR-D-TOPO previously (*25*), and 3×HA was cloned into pDONR P2r-P3. A F75G mutation was introduced into AFB1 CDS to make ccvAFB1 (*26*), similar to the ccvTIR1 published (*27*). Various GC and AC mutations were introduced into AFB1 CDS by using Phusion Site-Directed Mutagenesis Kit (Thermo Fisher, F541). The resulting entry clones were recombined into the destination vector pB7m24GW or pB7m34GW. All the primers used for plasmid construction are listed in table S1. The final expression constructs were transformed into the *Agrobacterium tumefaciens* strain GV3101 by electroporation. The floral dip method was used to transform the *Arabidopsis* plants.

Seeds were surface-sterilized by chlorine gas, sown on half-strength Murashige and Skoog (1/2 MS) medium supplemented with 1% (w/v) sucrose and 0.8% (w/v) phyto agar (pH 5.9), stratified in the dark at 4 °C for 2 days and then grown vertically at 21 °C with a long-day photoperiod (16 h light:8 h dark). The growth conditions are exactly same as previously described (*15*).

Protein purification

CDS sequences of TIR1ΔNT, AFB1 and AFB5 have been cloned into the pGEX-4T-1 vector before (*15*). The Phusion Site-Directed Mutagenesis Kit (Thermo Fisher, F541) was used to mutate the GC motifs in those plasmids using the primers listed in table S1. PaAFB1/2 and AmTrAFB1/2/3 coding sequences with AttB1/B2 sites were synthesized after codon optimization (Integrated DNA Technologies, USA), and were cloned into pDONR221. After confirmation with sequencing, they were amplified and re-cloned into the pGEX-4T-1 vector by the digestion-ligation approach. The primers used are listed in table S1. All the other constructs used for protein expression in this study have been published (*15*). The production and purification of the recombinant GST-tagged proteins from *E. coli* and the His-GFP-Flag-TIR1 protein from Sf9 insect cells, and excision of GST tags in some cases were performed following the procedures previously described (*15*).

*In vitro* GC and AC activity assays

*In vitro* GC activity of the purified proteins was determined by evaluating the rate of cGMP formation. The reaction mixture contained 10 mM Tris-HCl buffer (pH 7.6), 1 mM $MgCl_2$, 1 mM $MnCl_2$, 1 mM IBMX (3-isobutyl-1-methylxanthine), 1 mM GTP, 1 mM DTT and 5 µg of the protein in a final volume of 100 µL. To investigate the effects of IAA on the GC activities, 10 µM IAA was added to the reaction mixture along with 5 µg of IAA7 or IAA17. Samples were then incubated at 30 °C for 25 min. The enzyme reaction was terminated by incubation at 100 °C for 5 min and the samples were centrifuged at 16,100 g for 10 min. *In vitro* AC activity was performed exactly following the procedures described in the previous publication (*15*).

LC-MS/MS analysis

LC–MS/MS experiments were performed using the Nexera UHPLC and LCMS-8045 integrated system (Shimadzu). The ionization source parameters were optimized in positive ESI mode using pure cGMP dissolved in HPLC-grade water (Sigma) and the MS was operated in the positive-ion multiple reaction monitoring mode (MRM). Detection was based on selected reaction monitoring of cGMP by fragmenting its precursor ion at m/z 346.20 and yielding a product ion at m/z 152.25. Samples were separated at 40 °C using a XSelect CSH Phenyl-Hexyl column (100 × 2.1 mm, 3.5 µm, Waters). An isocratic flow of 90% solvent A (0.05% (v/v) formic acid with 5 mM ammonium formate) and 10% solvent B (100% (v/v) acetonitrile) was applied over 5 min, followed by washing and conditioning of the column, with a flow rate of 0.4 ml $min^{-1}$. The interface voltage was set at 4.0 kV for positive (ES+) electrospray. Data acquisition and analysis were made with the LabSolutions workstation for LCMS-8045.

Extraction and quantification of cAMP and cGMP in roots

Extraction of the total pool of cyclic nucleotides from roots was described in detail in the previous publication, and cAMP and cGMP were quantified in the same samples using UHPLC-MS/MS (*15*). For extraction and measurement of cGMP level in root tips by ELISA assay, a previously described method was used with minor modifications (*34*). Briefly, ~5-mm root tips were harvested from 5-day old seedlings and immediately frozen in liquid nitrogen. Approximately 400 mg of fresh ground powder was lysed in 500 µL of 5% TCA solution at 4 °C for 20 minutes. After centrifugation, supernatants were collected into new tubes and subjected to three sequential washes with 5 volumes of water-saturated ether. Samples were completely dehydrated and weighed to obtain the final dry biomass. Residual ether was removed by heating samples at 70 °C for 10 min. cGMP concentrations were determined using a cGMP ELISA kit (Cayman Chemical) following the standard protocol. Absorbance was measured with synergy H1 microplate reader (Biotek), and data were analyzed using the Cayman online tools (https://www.myassays.com/cyclic-amp-acetylated.assay).

Pull-down assays

*In vitro* pull-down assay to test the auxin-induced TIR1/AFBs-Aux/IAA interaction was done in a similar way as previously described (*15*). $TIR1^{GCms}$-HA and $ccvAFB1^{GCms}$-FLAG proteins were *in vitro* translated using TnT SP6 High-Yield Wheat Germ Protein Expression System (Promega, L3260), and GST-IAA7 protein was purified from *E. coli*. TIR1-HA in pF3A WG (BYDV) Flexi Vector (Promega, L5671) and GST-IAA7 in pGEX-4T-1 vector has been prepared before (*15*). The construct for ccvAFB1-FLAG was prepared in a similar way as TIR1-HA. Phusion Site-Directed Mutagenesis Kit (Thermo Fisher, F541) was used to generate the $TIR1^{GCms}$-HA and $ccvAFB1^{GCms}$-FLAG constructs using the primers listed in table S1. Pull-down reactions and western blotting were performed as previously described (*15*), using Glutathione agarose (Thermo Scientific, 16102) for pull-down, and anti-HA–peroxidase (Sigma, 12013819001), anti-FLAG-peroxidase (Sigma, H7425), anti-GST-tag, HRP-conjugated monoclonal antibody (Agrisera, AS18 4188) at a dilution of 1:2,000 for western blot.

Microfluidic vRootchip and live imaging

To evaluate auxin-induced rapid root growth inhibition and cytosolic $Ca^{2+}$ transients, the microfluidic vRootchip coupled to an in-house-established vertical Zeiss LSM 800 confocal microscope was used following the established procedures (*10, 12, 15, 35*). Simply, four-day-old seedlings were inserted into the channels of vRootchip with 1/4 MS liquid medium without sucrose supplied and were grown overnight. The vRootchip was then assembled into an imaging chamber and mounted onto the vertical microscope. Medium was changed to the basal medium

(1/4 MS + 0.1% sucrose) and the seedlings were adapted for more than 2 h before the start of imaging. Treatment medium was the same as the basal medium except containing 10 nM of IAA or 50 nM of cvxIAA. IAA was purchased from Sigma (I2886), and cvxIAA was from Tokyo Chemical Industry Co. (M3141). Both IAA and cvxIAA were dissolved in ethanol to prepare 20 mM stock, which was diluted into the medium to reach the final concentration. The calcium sensor GCaMP3 (*28*) was either crossed with different genotypes or being transformed with different constructs to monitor the cytosolic $Ca^{2+}$ level. Live imaging and data analysis were performed as previously described (*12, 15*). To evaluate multiples lines in a faster way, another simplified vRootchip system was also used as previously described (*13, 36*).

Measurement of $Ca^{2+}$ flux in root tips using microelectrodes

Roots from 7-day-old seedlings were immobilized on a glass slide with parafilm, and put in a Petri dish containing 30 mL BSM solution (Basic Salt Medium solution, 0.5 mM KCl, 0.1 mM $CaCl_2$) for 30 min. Calcium microelectrodes were prepared using Ca-selective ionophore I (99310, Sigma) and calibrated as described in our previous publications (*37*). Electrodes were positioned above the elongation zone of the root (300 µm from the root apex), and net $Ca^{2+}$ fluxes were measured using the MIFE non-invasive microelectrode ion flux measuring technique (*38*). The steady-state net $Ca^{2+}$ flux was first measured for 5 min, and then 10 nM IAA treatment was administered, followed by another 10-15 min of flux measurements. The magnitude of response (abbreviated as Δpeak flux) was calculated as the difference between initial (basal) value of flux and the peak value in response to IAA treatment for each replicate.

Electrical signal recording in root tips

Six-day-old seedlings were used for this experiment. A two-channel amplifier (Duo 773, World Precision Instruments, Friedberg, Germany) was used to record the root tip surface potential. To maintain a stable electrode potential, a glass pipette (2 cm long; inside diameter, 1.5 mm; outside diameter, 1.8 mm) was filled with 1/2 MS growth medium (which contains ~10 mM KCl). An Ag/AgCl wire electrode was inserted into the tube to the half-way point leaving the lower 1 cm part of the glass pipette containing 1/2 MS to form a junction with the agar on which the root tip was placed.

For experiments, seedlings were transferred to a new agar containing 1/2 MS growth medium. The end of the root protruded from the agar block and the root tip was placed on a small (3×3×3 mm) island block of the agar containing 1/2 MS growth medium. There was a 2-mm gap between the growth medium and the island block. The grounding/reference electrode was placed in the large agar block 5 cm from the seedlings. The recording electrode was placed close to the root on the island block to record the electrical signals. When a stable electrical signal baseline was established (few minutes after transferring plants), 5 µL IAA solution was applied to the root tip on the island block. Electrical signals were acquired in a Faraday cage at 100 Hz and analyzed using LabScribe4 software (iWorx Systems Inc., Dover, NH, USA).

External data sources

*Arabidopsis* gene and protein sequences are available from TAIR (https://www.arabidopsis.org/). TIR1/AFBs orthologues from *Picea abies* PaAFB1 (MA_14836g0010) and PaAFB2 (MA_121693g0010) were obtained from a BLAST search with the AtTIR1 protein sequence as bait from the Spruce Genome Project database (https://congenie.org/blast). The orthologous sequences from *Amborella trichopoda* AmTrAFB1 (v1.0_scaffold00016.85), AmTrAFB2 (v1.0_scaffold00003.426) and AmTrAFB3 (v1.0_scaffold00135.41) were identified from a

BLAST search in Phytozome 13 (https://phytozome-next.jgi.doe.gov/blast-search). The TIR1 3D structure (PDB 2P1Q) is available from PDBe (https://www.ebi.ac.uk/pdbe/entry/pdb/2P1Q).

Software and statistical analysis

Multiple sequence alignment was performed using the software Jalview (*39*). The TIR1 3D structure (PDB 2P1Q) was published (*40*), and was visualized and labelled in PDBe. All graphs were generated using GraphPad Prism 8. One-way ANOVA, two-way ANOVA and multiple comparisons were performed where necessary using GraphPad Prism 8.

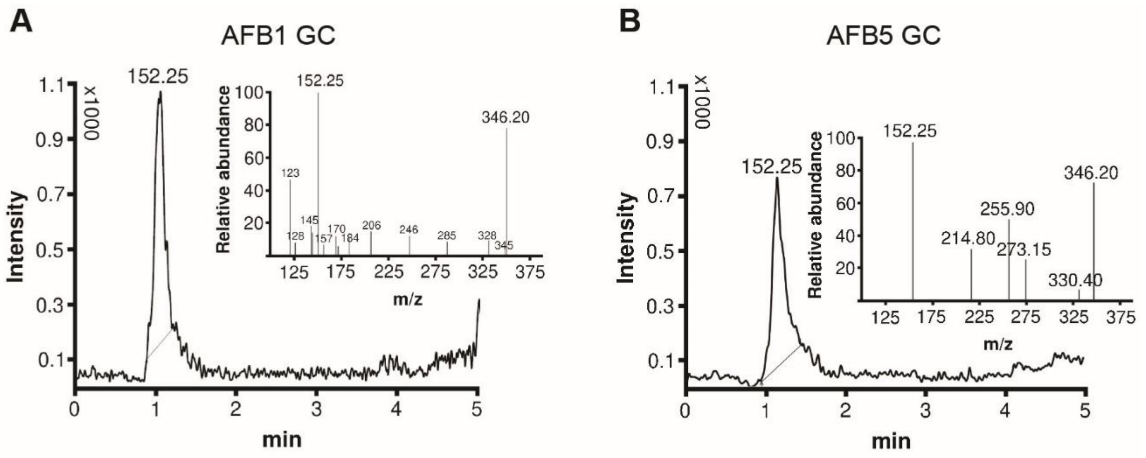


**Fig. S1. Additional data to support the GC activity of TIR1/AFBs.**

(**A**) LC-MS/MS spectrum for the GC reaction of GST-AFB1. (**B**) LC-MS/MS spectrum for the GC reaction of GST-AFB5. The typical MS/MS spectrum of cGMP was detected in the GC reactions with the characteristic chromatographic peak used for quantification.

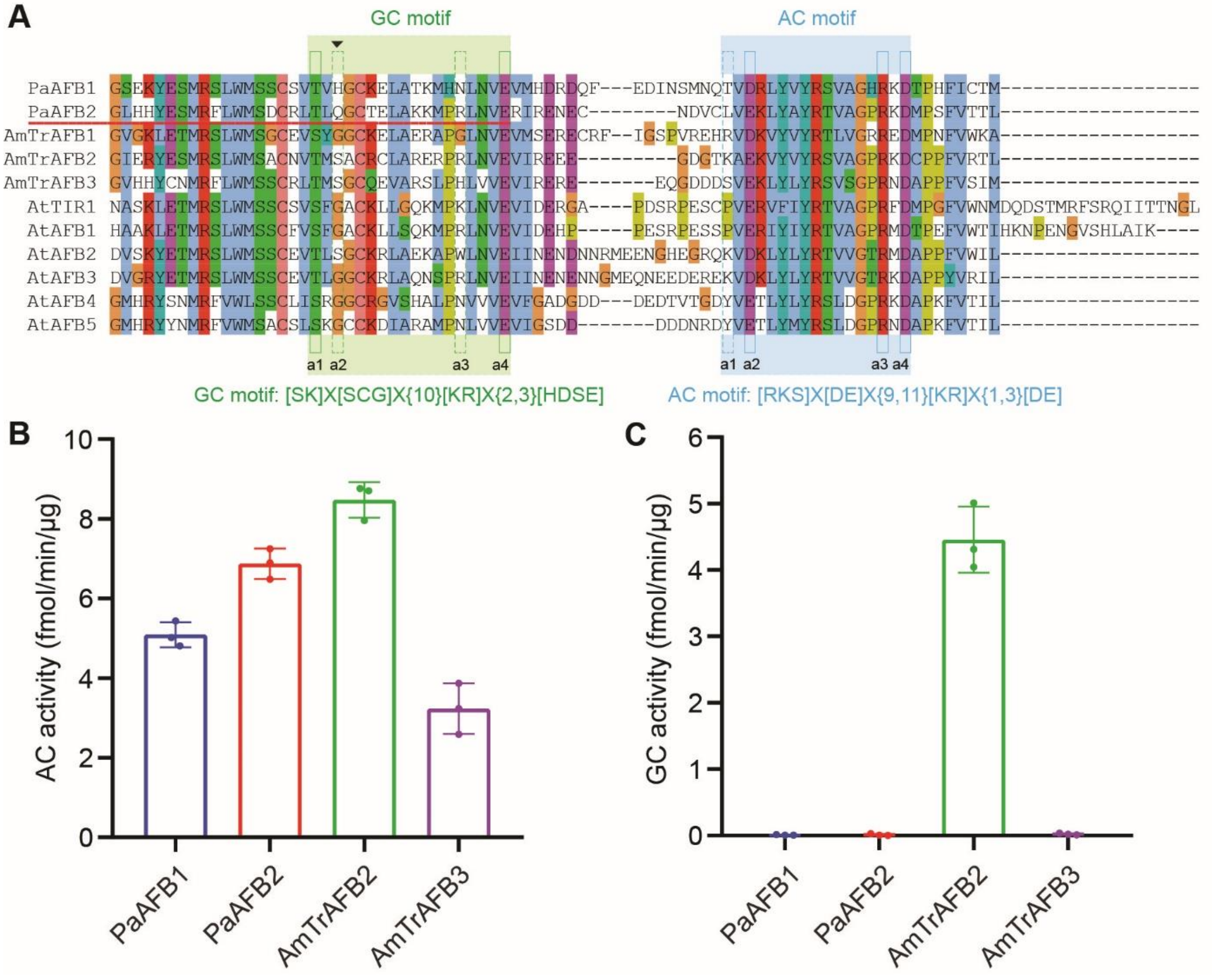


**Fig. S2. GC activity of TIR1/AFB orthologues evolved in angiosperms.**

(**A**) C-terminal regions of the multiple sequence alignments of TIR1/AFBs and their orthologues from the gymnosperm *Picea abies* (PaAFB1/2), and the ancestral angiosperm *Amborella trichopoda* (AmTrAFB1/2/3), showing the evolution of the GC motif. Note that there is a clear change of the 2nd key amino acid (a2) in GC motif between gymnosperm and angiosperm, indicated by a red line. In angiosperm, a2 of GC motif was evolved and fixed to perfectly match the reported GC motif. (**B**) All the TIR1/AFBs orthologues from *Picea abies* and *Amborella trichopoda* that can be purified have AC activity. (**C**) GC activity of TIR1/AFB orthologues evolved in angiosperm. GST-tagged proteins were purified from *E. coli*. *In vitro* AC/GC activity assay was performed, and cAMP/cGMP level after reaction was quantified using LC-MS/MS. Value from the GST negative control has been subtracted. The values shown are means ± s.d. from three biological replicates.

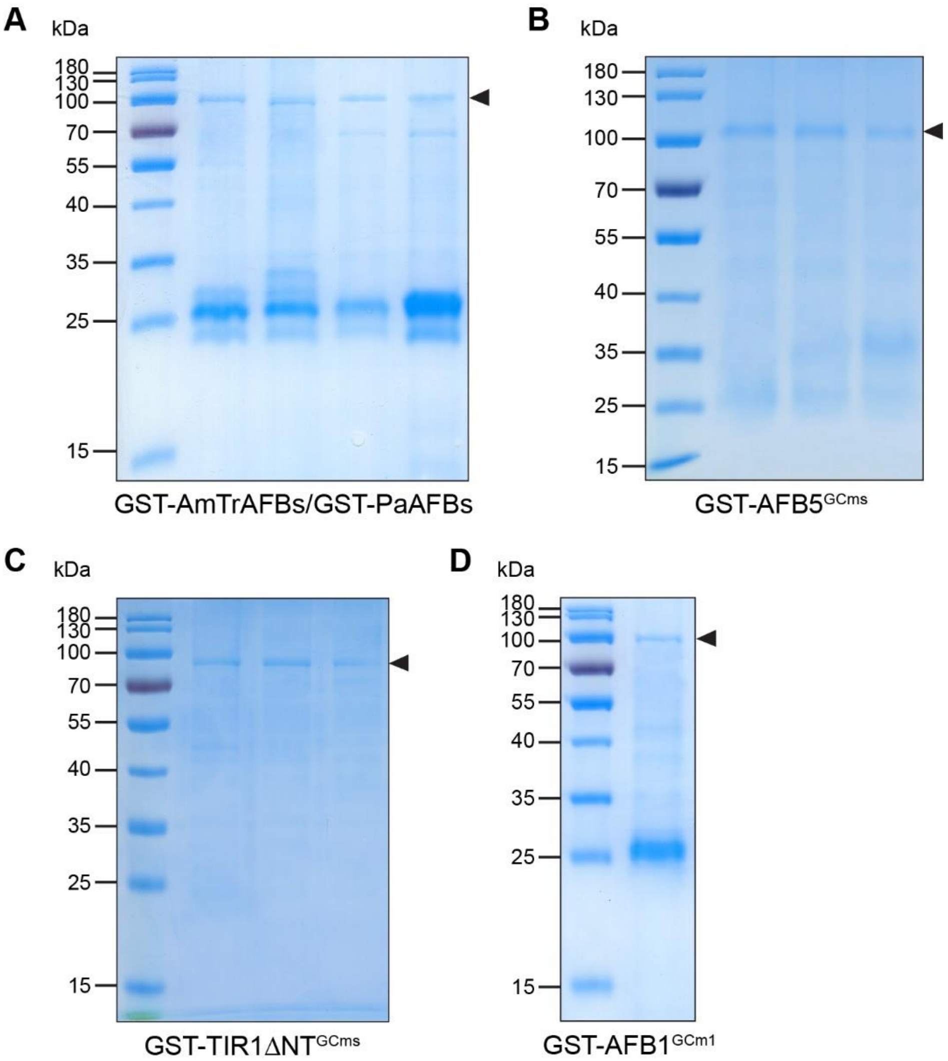


**Fig. S3. Gel images showing the purified proteins used in this study.**

(**A**) GST-AmTrAFB2/3, GST-PaAFB1/2. (**B**) GST-AFB5$^{GCm1/m2/m3}$. (**C**) GST-TIR1ΔNT $^{GCm1/m2/m3}$. (**D**) GST-AFB1$^{GCm1}$. All the proteins above were purified from BL-21 *E. coli* cells. Proteins were separated on SDS-PAGE gels and the gels were stained with Coomassie Brilliant Blue.

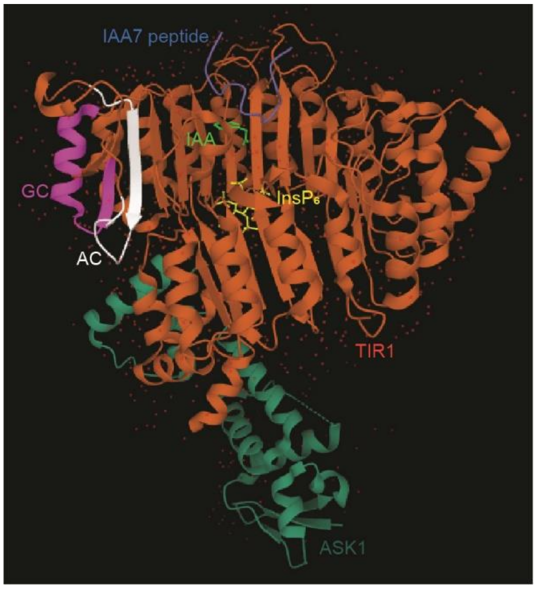


**Fig. S4. Protein structure of TIR1-IAA-Aux/IAA complex.**

Protein structure of TIR1-IAA-Aux/IAA complex showing the relative spatial position of the C-terminal GC and AC motifs. Different parts were labelled as different colors. Dark green, ASK1; Red, TIR1; Purple, C-terminal GC motif; White, C-terminal AC motif; Blue, IAA7 peptide; Yellow, InsP6 (inositol hexakisphosphate); Green, IAA.

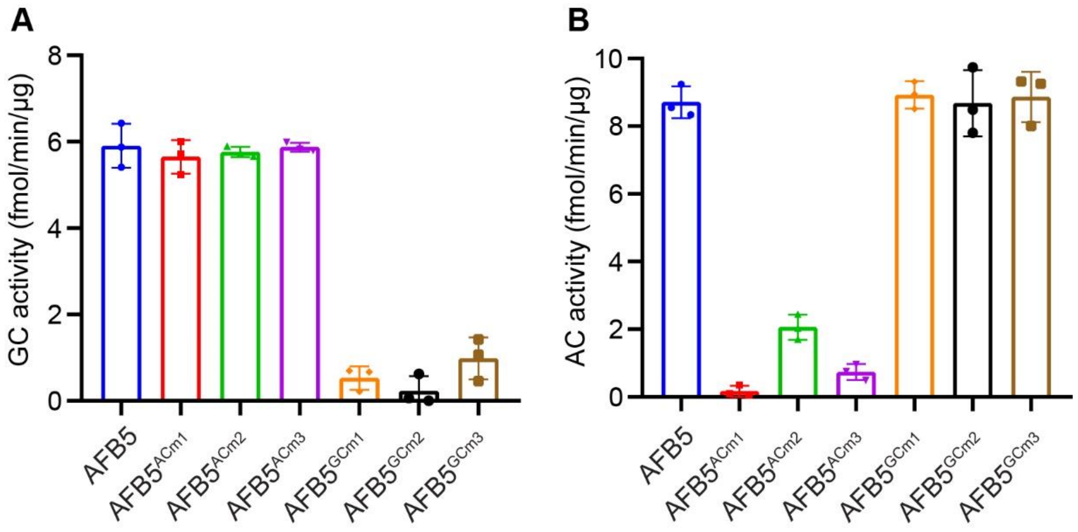


**Fig. S5. The C-terminal GC and AC motifs are specifically responsible for the GC or AC activity of AFB5.**

(**A**) The C-terminal GC motif but not the AC motif is specifically essential for the GC activity of AFB5. (**B**) The C-terminal AC motif but not the GC motif is specifically essential for the AC activity of AFB5. GST-AFB5, 3 AC motif mutated variants and 3 GC motif mutated variants were purified from *E. coli*. *In vitro* GC/AC activity assay was performed and cGMP/cAMP was quantified by LC-MS/MS. Data for the AC activity of AFB5 and AFB5$^{ACms}$ have been shown previously (*15*). Data are mean ± s.d. of three biological replicates.

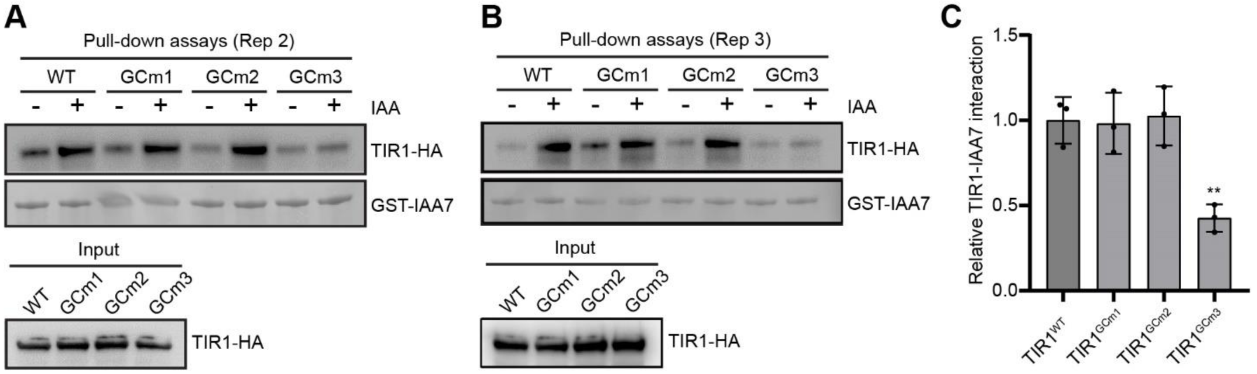


**Fig. S6. Pull-down assays showing differential effects of TIR1$^{GCms}$ mutations on the IAA-induced TIR1-Aux/IAA interaction.**

The TIR1-HA and its GCms variants were translated *in vitro* using wheat germ extracts. GST-IAA7 was purified from *E. coli*. Pull-down assays were performed in the presence or absence of 10 μM IAA as indicated. The experiments were repeated three times with similar results, and the two replicates were shown as the supplementary data to Fig. 2E (A and B). (C) Quantification of the TIR1-IAA7 interaction normalized to the input and the corresponding control without IAA. Data are mean ± s.d. of three biological replicates. One-way ANOVA and Dunnett's multiple comparisons test. ** $p \leqslant 0.01$.

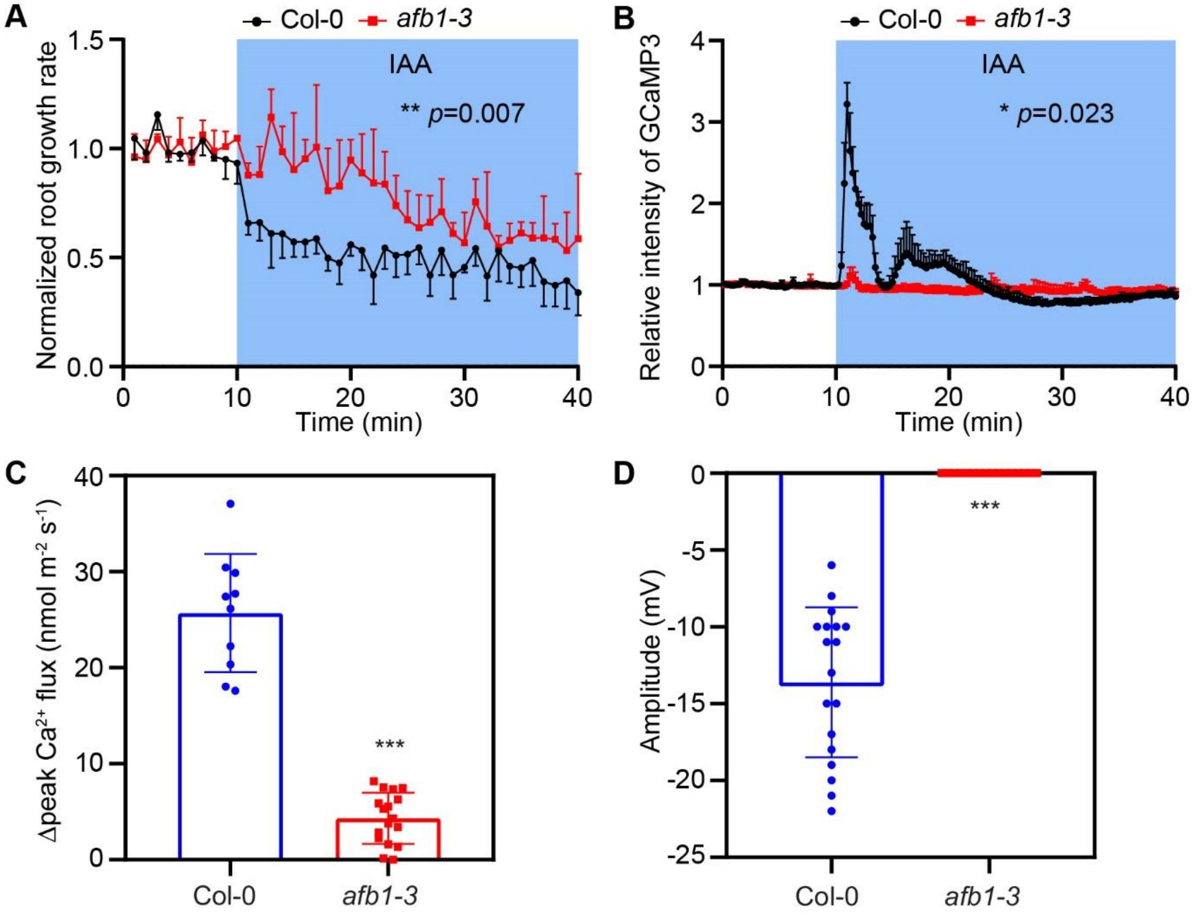


**Fig. S7. The *afb1* mutant is defective in rapid auxin responses.**

(**A**, **B**) AFB1 is required for IAA-induced rapid root growth inhibition (**A**) and cytosolic $Ca^{2+}$ transients (**B**). The vRootchip experiments were performed. Mock medium was changed to medium containing 10 nM IAA at 10 min. Normalized root growth rate and relative fluorescence intensity of GCaMP3 in elongation zone were quantified. Data shown are means +/- s.d. n = 4 seedlings. Two-way ANOVA and Sidak's multiple comparisons test. The *p*-values for genotypic variance are shown. (**C**) Auxin-induced $Ca^{2+}$ flux in root tips. Seven-day-old seedlings were used for analysis. The difference of $Ca^{2+}$ flux between basal values and peak values in response to 10 nM IAA was measured by $Ca^{2+}$-selective microelectrodes. Data are mean ± s.d. n = 10-17 measurements from 6-10 seedlings. Student's t-test, *** $p \leq 0.001$. (**D**) Auxin-induced electrical signal in root tips. Six-day-old seedlings were used for analysis. Electrical signals in root tips after treatment with 100 nM IAA were recorded. Data are mean ± s.d. n = 16-18 seedlings. Student's t-test, *** $p \leq 0.001$.

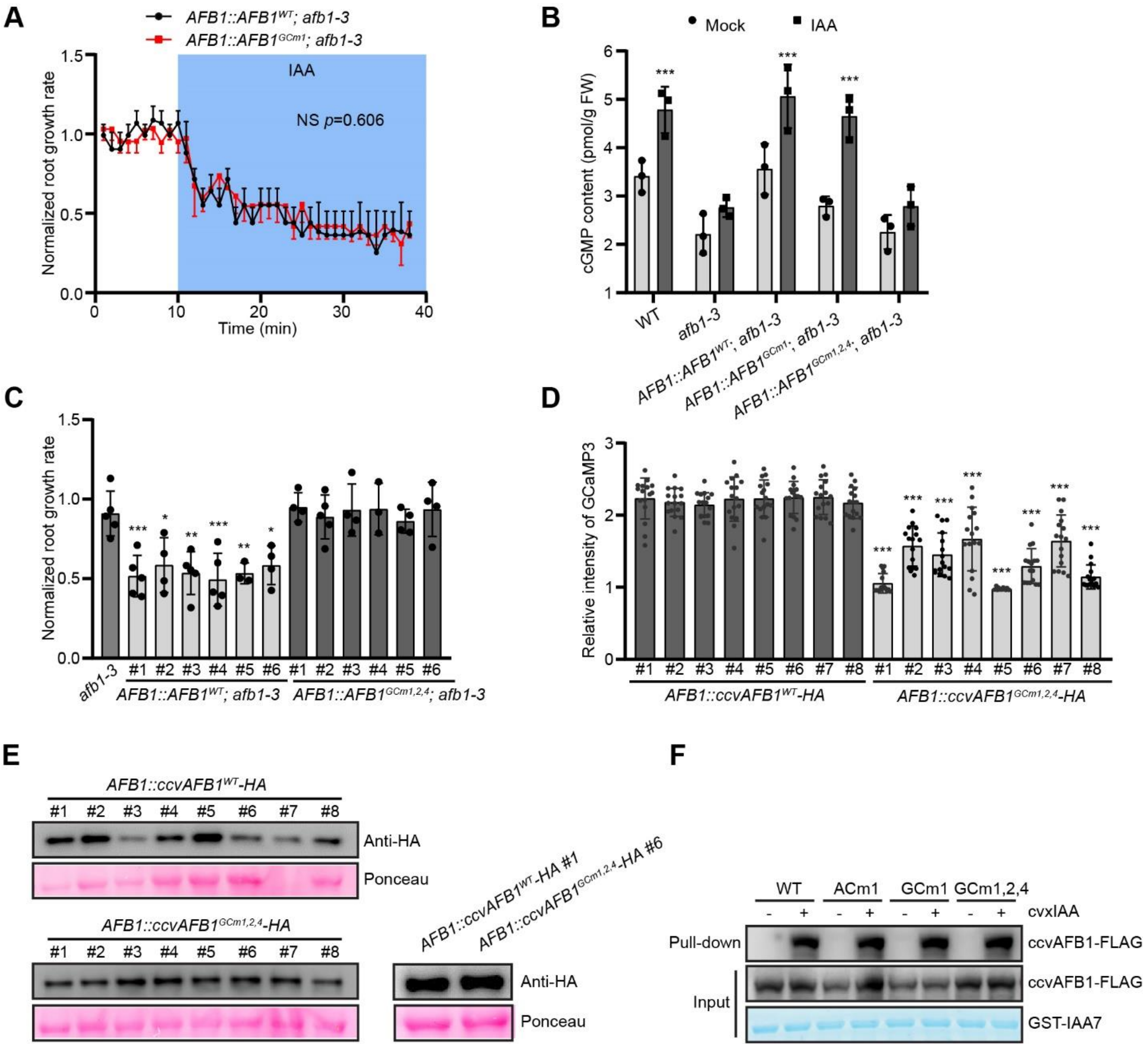


**Fig. S8. The GCm1,2,4 mutation in AFB1 specifically abolishes its GC activity and largely compromises its ability to mediate rapid auxin responses.**

(**A**) AFB1$^{GCm1}$ complements the *afb1-3* mutant normally as AFB1$^{WT}$ in terms of IAA-induced rapid root growth inhibition. The vRootchip experiments were performed. Mock medium was changed to medium containing 10 nM IAA at 10 min. Normalized root growth rate was quantified. Data shown are means +/- s.d. n = 3 seedlings. Two-way ANOVA and Sidak's multiple comparisons test. The *p*-values for genotypic variance are shown. (**B**) AFB1$^{GCm1,2,4}$ mutation largely abolishes the auxin-induced cGMP accumulation in root tips as determined by ELISA, while AFB1$^{GCm1}$ does not. Five-day-old seedlings of the indicated genotypes were treated with Mock or 100 nM IAA for 1 min, and approximately 5-mm root tips were harvested for cGMP quantification by an ELISA assay. Data are mean ± s.d. from three biological replicates. Two-way ANOVA and Tukey's multiple comparisons test. Statistical significance between Mock and IAA group for each genotype is shown. *** $p \leqslant 0.001$. (**C**) AFB1$^{GCm1,2,4}$ failed to complement the *afb1-3* mutant in terms of IAA-induced rapid root growth inhibition in

6 independent lines. The vRootchip experiments were performed with the indicated genotypes. Shown are mean ± s.d. of the normalized root growth rate at 2 min after treatment with 10 nM IAA. n = 3-5 seedlings. One-way ANOVA and Dunnett's multiple comparisons test. Statistical significance for each line compared to *afb1-3* is shown. * $p \leqslant 0.05$; ** $p \leqslant 0.01$; *** $p \leqslant 0.001$. (**D**) cvxIAA-induced $Ca^{2+}$ spike is largely reduced in 8 independent *ccvAFB1*$^{GCm1,2,4}$*-HA* lines compared to the *ccvAFB1*$^{WT}$*-HA lines*. The vRootchip experiments were performed with the indicated lines transformed into GCaMP3 background. Relative fluorescence intensity of GCaMP3 in elongation zone were quantified at 3 min after treatment with 50 nM cvxIAA. Shown are mean ± s.d. n = 16 seedlings. One-way ANOVA and Dunnett's multiple comparisons test. Statistical significance compared to the first line is shown. *** $p \leqslant 0.001$. (**E**) The *ccvAFB1*$^{WT}$*-HA* and *ccvAFB1*$^{GCm1,2,4}$*-HA* lines have comparable protein expression level. Western blot was performed with 6-day-old seedlings. Ponceau S stain serves as loading control. (**F**) The GCm1,2,4 mutation in ccvAFB1 does not affect its interaction with IAA7 in the presence of cvxIAA. The ccvAFB1-FLAG and its variants were translated *in vitro* using wheat germ extracts. GST-IAA7 was purified from *E. coli*. Pull-down assays were performed in the presence or absence of 10 μM cvxIAA as indicated.

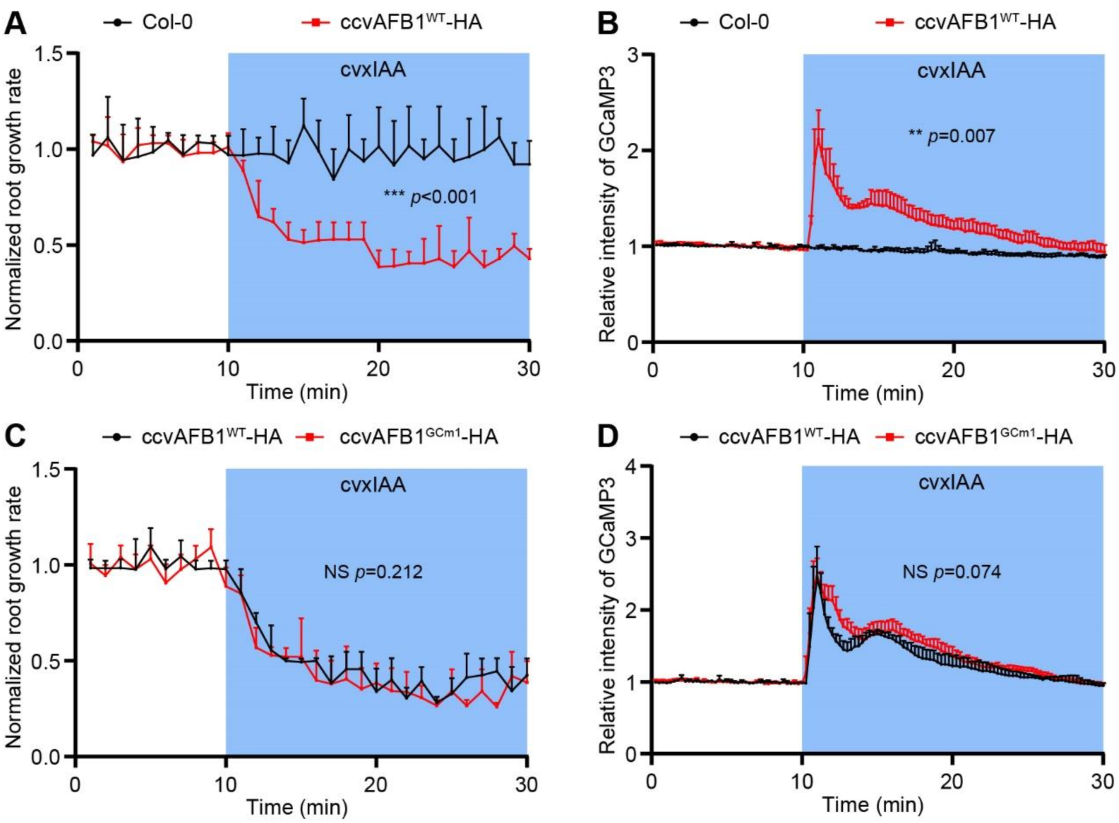


**Fig. S9. The GCm1 mutation in ccvAFB1 does not affect cvxIAA-induced rapid root growth inhibition and $Ca^{2+}$ spikes.**

Wild type, *AFB1::ccvAFB1$^{WT}$-HA* and *AFB1::ccvAFB1$^{GCm1}$-HA* were respectively crossed with GCaMP3, and the 5-day-old F1 seedlings were directly used for vRootchip experiment. Mock medium was changed to medium containing 50 nM of cvxIAA at 10 min. Normalized root growth rate (A and C) and Relative intensity of GCaMP3 signal (B and D) in epidermal cells of elongation zone were measured. The values were normalized to the average before treatment of the respective group. Shown are means + s.d. n = 4 seedlings. Two-way ANOVA and Sidak's multiple comparisons test. The *p*-values for genotypic variance are shown.

**Table S1. Primes used in this study.**

| Primers | Sequences | Usage |
|---|---|---|
| *afb1-3*-GT-FP | AACGGAAGACTAGGAAGCGAG | Genotyping |
| *afb1-3*-GT-RP | GCAACAGCTTCAAGACCTTTG | |
| LBb1.3 | ATTTTGCCGATTTCGGAAC (for *afb1-3*) | |
| PaAFB1-pGEX-FP | CCGGAATTCATGGTTGTCCGCCGGATGTCATC (EcoRI) | Protein expression in *E. coli* |
| PaAFB1-pGEX-RP | ATAAGAATGCGGCCGCTTACATTGTACAGATAAAGTGTGGTGTGTC (NotI) | |
| PaAFB2-pGEX-FP | CCGGAATTCATGGATCAGGTGCAATCTATTTGCC (EcoRI) | |
| PaAFB2-pGEX-RP | ATAAGAATGCGGCCGCCTAAAGCGTTGTAACGAAGGAAGGC (NotI) | |
| AmTrAFB1-pGEX-FP | CCGGAATTCATGACTTCGTTTCCTGAAGAGGTATTGG (EcoRI) | |
| AmTrAFB1-pGEX-RP | ATAAGAATGCGGCCGCCTAAGCCTTCCATACAAAGTTGGGC (NotI) | |
| AmTrAFB2-pGEX-FP | CCGGAATTCATGGGTCTTCGGAGCGAAAAACG (EcoRI) | |
| AmTrAFB2-pGEX-RP | ATAAGAATGCGGCCGCTCACAAAGTTCTTACGAAGGGCG (NotI) | |
| AmTrAFB3-pGEX-FP | CCGGAATTCATGACCCATGACCAACCCTCTGC (EcoRI) | |
| AmTrAFB3-pGEX-RP | ATAAGAATGCGGCCGCTTACATTATTGATACAAATGGCGGTGCA (NotI) | |
| AFB1p-B4-FP | CAACTTTGTATAGAAAAGTTGGGCATGGAGAACATAAACGAATCAACTA | Transgenic plants |
| AFB1p-B1r-RP | CGTTCAACTTTTTTGTACAAACTTGACAGAAACAGAGTGAGAGAGAGAG | |
| AFB1-CDS-FP | GGCCGCCCCCTTCACCATGGGTCTCCGATTCCCACCTAA | |
| AFB1-CDS-RP | CGGCGCGCCCACCCTTCTTTATGGCTAGATGTGAAACTCCAT | |
| AFB1-ccv-FP | GGAAGCCACACGGCGCTGACTATAATC | ccvAFB1 mutation |
| AFB1-ccv-RP | CCTTCAAAGTCAAAGATCTCATCTCCGG | |

| | | |
|---|---|---|
| TIR1-GCm1-FP | GTGAGTTTTGCAGCCTGCAAGTTAC | TIR1 GC motif mutations |
| TIR1-GCm1-RP | GGAACAAGAAGACATCCAAAGGGATC | |
| TIR1-GCm2-FP | CAGAAGATGCCAGCGCTGAATGTG | |
| TIR1-GCm2-RP | TCCTAGTAACTTGCAGGCTCCAAAAC | |
| TIR1-GCm3-FP | GCTGAATGTGGCAGTCATCGATGAAC | |
| TIR1-GCm3-RP | TTTGGCATCTTCTGTCCTAGTAACTTG | |
| AFB5-GCm1-FP | TAGCTTGTCTAAGGCATGCTGCAAGGATATTG | AFB5 GC motif mutations |
| AFB5-GCm1-RP | CATGCTGACATCCAAACAAACCTCATGTTGTAATAG | |
| AFB5-GCm2-FP | GAGCAATGCCGGCTCTAGTTGTGGAAG | |
| AFB5-GCm2-RP | GTGCAATATCCTTGCAGCATCCCTTAGAC | |
| AFB5-GCm3-FP | CGAATCTAGTTGTGGCAGTAATTGGATCGGATG | |
| AFB5-GCm3-RP | GCATTGCTCGTGCAATATCCTTGCAG | |
| AFB1-GCm1-FP | CTTTGTAAGTTTTGCTGCTTGCAAGCTTC | AFB1 mutations |
| AFB1-GCm1-RP | CAAGATGACATCCAAAGGGATCGCATG | |
| AFB1-GCm2-FP | CAGAAAATGCCAGCGCTCAATGTCGAAG | |
| AFB1-GCm2-RP | ACTTAGAAGCTTGCAAGCACCAAAACTTAC | |
| AFB1-GCm4-FP | AGTCAGAAAATGGCAAGGCTCAATGTCG | |
| AFB1-GCm4-RP | TAGAAGCTTGCAAGCACCAAAACTTACA | |
| AFB1-ACm1-FP | GAGCTCTCCAGTTGCGAGGATATACATATAC | |
| AFB1-ACm1-RP | TCAGGTCTTGACTCTGGAGGATGTTC | |
| pGEX-S-F | GGGCTGGCAAGCCACGTTTGGTG | Sequencing |
| pGEX-S-R | CCGGGAGCTGCATGTGTCAGAGG | |
| M13-21F | TGTAAAACGACGGCCAGT | |
| M13-29R | CAGGAAACAGCTATGACC | |
